# Exploiting Homeostatic Repopulation to Increase DC Vaccine Efficacy in Multiple Myeloma

**DOI:** 10.1101/049072

**Authors:** Chandler D. Gatenbee, Núria Folguera-Blasco, Charlie Daneils, Jill Gallaher, Russell C. Rockne, Casey Adams, Michael Nicholson, Eleni Maniati, John Kennedy, Kimberly Luddy, Frederick L. Locke, Mark Robertson-Tessi

## Abstract

Standard of care for multiple myeloma involves autologous hematopoietic cell transplant (AHCT), which can extend life by a year; however the disease remains incurable. A dendritic cell vaccine developed at Moffitt will be used both immediately before and after AHCT, with the aim of achieving complete response. Data will be collected during the trial to test the biological activity of the vaccine. This data will parameterize a model that facilitates the exploration of outcomes when varying the timing of vaccination. This calibrated model will also inform the design of a follow-up trial, which will include vaccination in conjunction with other immunotherapies.

## I BIOLOGICAL BACKGROUND

### Ia Multiple Myeloma

Multiple myeloma is the second most common hematologic malignancy in adults, with approximately 26,850 patients diagnosed per year in the United States [10]. The disease is characterized by the proliferation of clonal plasma cells preferentially in the bone marrow, resulting in anemia, osteolytic bone disease and the detection of a monoclonal gammopathy in the majority of the patients [3]. The current standard therapy consists of induction therapy with immunomodulatory drugs or proteasome-inhibitor-based regimens, followed by autologous hematopoietic cell transplants (AHCT) in those patients with responsive disease [2, 7, 8, 9]. These treatment modalities induce high rates of complete remission and have significantly improved survival. However, molecular remissions are rare, and a significant proportion of patients are unable to achieve a complete response (CR) to induction therapy, both before and after transplant. Inevitably all patients relapse and die due to disease progression [6]. Therefore, the development of novel interventions for patients with resistant disease, and the targeting of minimal residual disease after autologous bone marrow transplantation, is greatly needed.

### Ib Clinical Trial for Dendritic Cell Vaccine Against Survivin

The period of T-cell homeostatic repopulation following AHCT provides a unique opportunity to stimulate a potent immune response against residual tumor cells. Tumors are often able to resist the immune responses by the induction of peripheral tolerance in the T-cell population. However, it has been shown that the T-cell repopulation phase post T-cell depletion results in a break from peripheral tolerance, opening up the opportunity for tumor-antigen-specific T-cell clones to expand in an effort to eradicate the tumor. The protein survivin functions as an apoptosis inhibitor via spindle microtubule and mitotic checkpoint regulation [1, 5]. It is expressed in development, undetectable in most adult tissues [11], yet overexpressed in almost all cancers [2, 4, 12, 13]. Such properties make survivin an excellent candidate as a tumor associated antigen that can be targeted by dendritic cell (DC) vaccines. In this vein, a novel fulllength DC vaccine against survivin has been developed, designed to maximize both immunogenicity and the number of patients eligible for treatment. It is our hypothesis that administration of this vaccine during homeostatic repopulation following AHCT will induce the clonal expansion of cytotoxic T-cells specific to survivin-positive tumor cells, producing a robust, therapeutic immune response against myeloma. We are currently preparing to undertake an approved clinical trial to test the safety and biological activity of this DC survivin vaccine. During this trial we will monitor the tumor and immune response to the vaccine over 6 months, taking numerous measurements at five time points: before transplantation, after stem cell mobilization/collection, and then 60, 90, and 180 days following AHCT

## II MATHEMATICAL MODEL

### IIa Motivation

We have developed a mathematical model that utilizes the wealth of data that will be gathered during this clinical trial. The model will be parameterized using this data, and will then be used to inform the design of a follow-up trial, which will include additional immunotherapies. The goal of this model is thus twofold: 1) Determine the optimum time at which to administer the DC vaccine, so as to maximize the expansion of cytotoxic T-cells specific for survivin positive tumor cells; 2) Explore the impact of administering additional immunotherapies, such as anti-PD-L1 or anti-CTLA4, which may augment the efficacy of the survivin vaccine.

The immune system is a complex mix of many cell types. To be included in the model, cell types had to meet certain criteria. First, the interactions of the cells selected must be well characterized. Second, the number of each cell type must be clinically measureable. Third, the interactions must either play a direct role in homeostatic repopulation and/or be targetable by available therapies. Finally, all cells and interactions must come together to form a model capable of realistically recapitulating the dynamics of homeostatic re-population. The requirement that the size of each cell population be measureable will be particularly important when simulating tumor growth and treatment using individual patient data. Fortunately, we have access to a number of methods that allow us to do just that: flow cytometry will be used to measure the number of regulatory T-cells and the overall T-cell category; nanostring measurements will determine the number of survivin-positive effector T-cells, exhausted T-cells, and activated T-cells; and M protein and light chain ratios will measure the number of tumor cells. Having data at multiple time points also gives us the ability accurately estimate many of the parameters in our model.

### IIb Equations

We have built a preliminary mathematical model consisting of a set of ordinary differential equations (ODEs) to describe the tumor burden and the immune response. The current model is developed for the period of time immediately following transplantion of stem and immune cells in the clinical protocol, such that the T-cells and tumor cells are both repopulating the depleted microenvironment. The system of 6 ODEs considers the following cell populations: tumor burden (*P*); activated T-cells (*A*); effector T-cells (*E*); tolerized T-cells (*X*); regulatory T-cells (*R*); and overall T-cells (*T*). The following system of ODEs was used as a preliminary model of the dendritic cell therapy.

1. 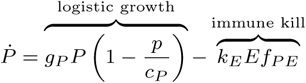
2. 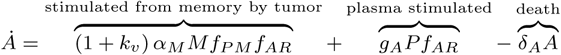
3. 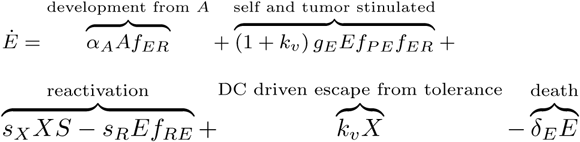
4. 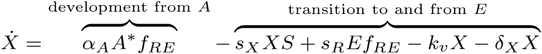
5. 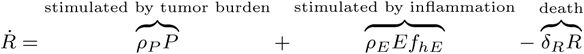
6. 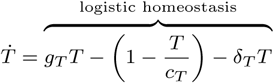

Memory cells (*M*) are a constant fraction of the total T-cell population (*T*). The variable *S* is the available space in the T cell compartment, calculated as 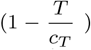 The final term of Eqs. 2-6 represents the natural death of that compartment. The term (1+ *kv*) represents the effect of the dendritic cell therapy. In all equations, the following shorthand is used:

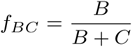

The tumor burden (Eq. 1) is increased by logistic growth (first term) and decreased through killing by the effector T-cells (second term). Activated T-cells (Eq. 2) are stimulated (first term) from memory cells by increased tumor burden; high Treg numbers suppress their stimulation and self proliferation (first and second term). Effector T-cells (Eq. 3) develop from the activated T-cells (first term) and through self proliferation based on tumor burden (second term). The third and fourth terms represent transition to and from the tolerized state, affected by space and Treg numbers. Dendritic cell therapy enhances the escape from tolerance (fifth term).

Tolerized T-cells (Eq. 4) are derived from activated T-cells when Treg numbers are high (first term). Terms 2-4 are the transition to and from the effector T-cell compartment. Tregs (Eq. 5) are promoted by the tumor burden (first term) and the amount of inflammation in the system (second term). The total T-cell compartment (Eq. 6) is modeled as logistic homeostasis. See Figure 1 for a schematic illustrating these interactions.

**Figure 1:**
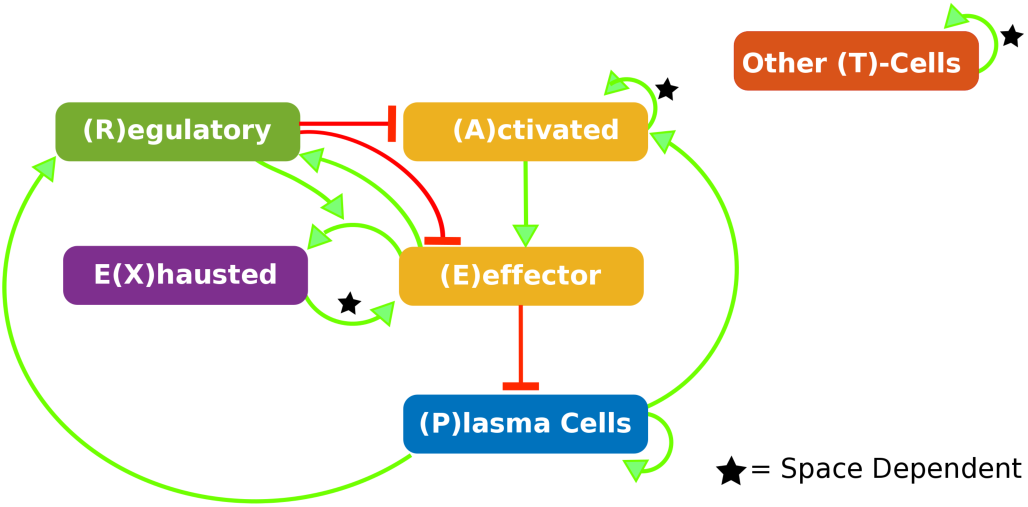
Interaction Network. See Equations section for full description.

## III ILLUSTRATIVE RESULTS OF APPLICATION OF METHODS

A sample result of our model’s ability to simulate the effect of timing of the vaccine can be found in Figure 2. These simulations highlight the importance timing has on the ability of the vaccine to stimulate a sufficient anti-tumor immune response. In panels A and C, the vaccine was given too early (day 10) and too late (90), respectively. In these cases, treatment fails because it misses the window of opportunity in which homeostatic repopulation can be exploited to increase expansion of survivin specific cytotoxic T-cells. Giving the vaccine too early will fail because there are an insufficient number of T-cells to stimulate, while administering it too late it will fail due to the return of peripheral tolerance. However, there is a sweet spot, where the T-cell population is neither too large nor too small, during which the vaccine will stimulate an immune response strong enough to eliminate the tumor (panel D). In these simulations, this ideal time to administer the vaccine occurs 50 days post AHCT (panel B). The optimum is sensitive to the parameters and initial conditions, suggesting that the model can be used in a patient-specific way to predict individual optima for vaccination.

**Figure 2:**
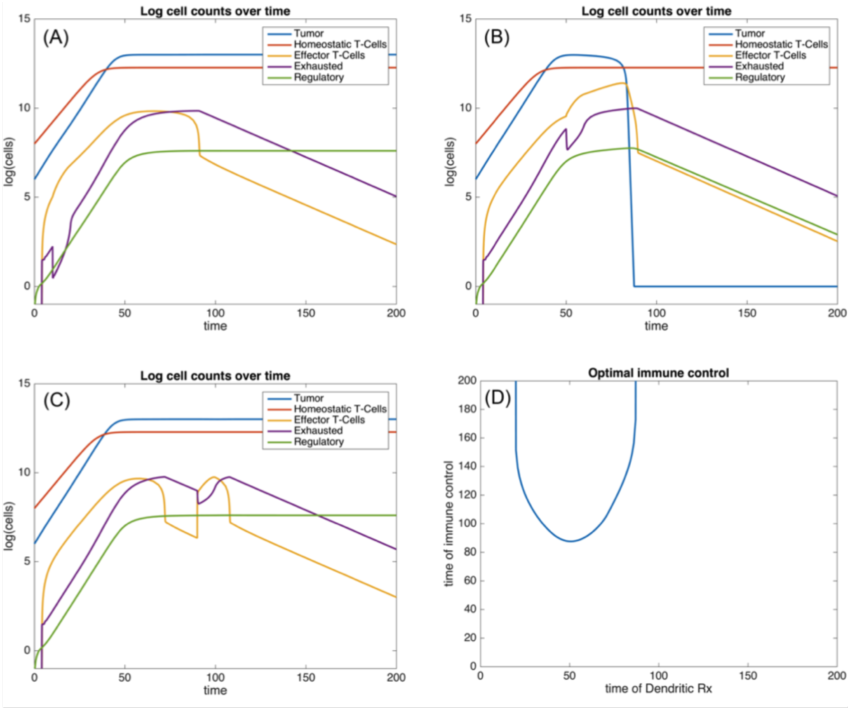
Results from the model showing application of dendritic cell therapy at time points 10, 50, and 90 days post transplant (panels A-C respectively). Only the therapy at 50 days leads to immune control of the tumor. Panel D shows the optimum tumor control as a function of the day of the dendritic cell therapy application. The vertical axis measures the time at which the tumor reaches its minimum post-dendritic therapy level.

## IV CLOSING REMARKS

The nature of multiple myeloma and the ability to collect key data at multiple time points provides the unique opportunity to develop a highly parameterized model of homeostatic T-cell repopulation, tumor growth, and the efficacy of immunotherapy. It is our hope that the model developed here will serve as a powerful tool for clinicians designing future clinical trials.

## ACKNOWLEDGEMENTS

We would like to thank Dr. Alexander R. A. Anderson and the Moffitt Cancer Center for organization and support of the 5th Annual Integrated Mathematical Oncology workshop, Immune Cancer, where this project was conceived. N.F-B. acknowledges MINECO for funding under grant MTM2015-71509-C2-1-R and Generalitat de Catalunya for funding under grant 2014SGR1307. N.F-B. is supported by a grant of the Obra Social La Caixa Foundation on *Collaborative Mathematics* awarded to the Centre de Recerca Matemàtica.

## References

[1] DC Altieri. Survivin, versatile modulation of cell division and apoptosis in cancer. Oncogene, 22(53):8581–8589, 2003.

[2] G Ambrosini, C Adida, and DC Altieri. A novel anti-apoptosis gene, survivin, expressed in cancer and lymphoma. Nat Med., 3(8):917–21, 1997.

[3] B Barlogie, J Shaughnessy, and G Tricot. Treatment of multiple myeloma. Blood, 103:20–32, 2004.

[4] H Kawasaki, DC Altieri, CD Lu, M Toyoda, T Tenjo, and N Tanigawa. Inhibition of apoptosis by survivin predicts shorter survival rates in colorectal cancer. Cancer Res., 58(22):5071–5074, 1998.

[5] SK Knauer, W Mann, and Stauber RH. Survivin’s dual role: An export’s view. Cell Cycle, 5(5):518–21, 2007.

[6] Y Nakagawa, S Abe, and Kurata M. Iap family protein expression correlates with poor outcome of multiple myeloma patients in association with chemotherapy-induced overexpression of multidrug resistance genes. Am J Hematol, 81:824–31, 2006.

[7] A Palumbo, JS Miguel, and Sonneveld P. Lenalidomide: A new therapy for multiple myeloma. Cancer Treat Rev, 34:283–91, 2008.

[8] AC Rawstron. Minimal residual disease detection in myeloma: No more molecular remissions? Haematologica, 90, 2005.

[9] Schlossman R Richardson PG, Mitsiades C. Bortezomib in the front-line treatment of multiple myeloma. Expert Rev Anticancer Ther, 8:1053–72, 2008.

[10] American Cancer Society. Cancer facts and figures 2016. Atlanta, GA, 2016.

[11] RH Stauber, W Mann, and SK Knauer. Nuclear and cytoplasmic survivin: Molecular mechanism, prognostic, and therapeutic potential. Cancer Res, 67(13):5999–6002, 2007.

[12] HS Swana, D Grossman, JN Anthony, RM Weiss, and Altieri DC. Tumor content of the antiapoptosis molecule survivin and recurrence of bladder cancer. N Engl J Med, 341(6):452–53, 1999.

[13] K Tanaka, S Iwamoto, G Gon, T Nohara, M Iwamoto, and Tanigawa N. Expression of survivin and its relationship to loss of apoptosis in breast carcinoma. Clin Cancer Res, 6(1):127–134, 2000.

